# Hidden potential of the supporting scaffold as a structural module for plant cystatin design

**DOI:** 10.1101/2022.02.26.482125

**Authors:** Jonathan Tremblay, Marie-Claire Goulet, Charles Goulet, Dominique Michaud

## Abstract

Protein engineering approaches have been proposed to improve the inhibitory properties of plant cystatins towards herbivorous pest digestive Cys proteases, typically involving sequence alterations in the inhibitory loops and/or N-terminal trunk of the protein interacting with specific amino acid residues of the target protease. In this study, we assessed whether the loops-supporting frame, or scaffold, would represent a valuable structural module for cystatin function improvement. Twenty hybrid cystatins were designed *in silico*, consisting of the N-terminal trunk and two inhibitory loops of a given donor cystatin grafted onto the scaffold of an alternative, recipient cystatin. Synthetic genes for the hybrids were expressed in *E. coli*, and the resulting proteins assessed for their potency to inhibit model Cys protease papain and the digestive Cys proteases of Colorado potato beetle (*Leptinotarsa decemlineata*) used as an insect pest model. In line with the occurrence of positively selected amino acids presumably influencing inhibitory activity in the scaffold region of plant cystatins, grafting the N-terminal trunk and inhibitory loops of a given cystatin onto the scaffold of an alternative cystatin generally had an effect on the inhibitory potency of these function-related elements against Cys proteases. For instance, hybrid cystatins including the three structural elements of model tomato cystatin SlCYS8 grafted on the scaffold of cystatins from other plant families showed *K*_i_ values altered by up to 3-fold for papain, and inhibitory efficiencies increased by up to 8-fold against *L. decemlineata* cathepsin L-like proteases, compared to wild-type SlCYS8 bearing the original scaffold. Our data point to a significant influence of the cystatin scaffold on the inhibitory activity of the N-terminal trunk and protease inhibitory loops. They also suggest the potential of this structural element as a module for plant cystatin design to generate functional variability against Cys proteases, including the digestive proteases of herbivorous pests.

## 1 INTRODUCTION

Numerous studies have discussed the protective effects of plant cystatins against herbivorous arthropods.^1,2^ These proteins act as pseudosubstrate inhibitors to obstruct the active site cleft of Cys proteases and block their catalytic activity on protein substrates.^3^ Upon feeding, cystatins inhibit digestive proteases in the arthropod’s digestive tract, to cause amino acid shortage leading to growth delays, developmental defects, and eventual death of the herbivore.^4^ From a physiological standpoint, the plant protective effect of a given cystatin is determined by its relative abundance compared to target Cys proteases in the arthropod midgut, by its basic inhibitory potency against these enzymes, and by eventual compensatory responses triggered in the herbivore following plant tissue intake.^5^ Herbivorous arthropods have developed effective strategies to cope with dietary protease inhibitors, notably involving the production of protease isoforms weakly sensitive to inhibition and the overexpression of protease-sensitive proteases to compensate for the loss of protein digestive functions.^6^

A key challenge to improve the protective effects of plant cystatins is to identify, or to develop, cystatin variants that show improved potency and/or a broadened inhibitory range against the target arthropod digestive proteases.^2,7^ Different strategies have been proposed for the molecular improvement of cystatins towards Cys proteases, usually involving primary sequence mutations in function-related regions of the protein. The inhibitory function of plant cystatins relies on two hairpin loops that establish physical interactions with specific amino acid residues in the active site cleft of the target enzyme.^8^ The N-terminal region, or N-terminal trunk, which interacts with surface residues on the target enzyme, also has a strong influence on the inhibitory potency and specificity of the cystatin towards different protease isoforms, owing to a steric impact on the spatial orientation of the inhibitory loops.^9,10^ Accordingly, protein engineering schemes involving site-directed mutagenesis or random mutations of functionally relevant amino acids in the N-terminal trunk or inhibitory loops of plant cystatins allowed for the design of improved cystatin variants potentially useful in plant protection against herbivorous pests or pathogens.^2,11^ Likewise, a newly described strategy for protein design that involves the substitution of function-related structural elements by the corresponding elements of an alternative cystatin allowed for the design of cystatin variants with highly improved inhibitory activity against digestive Cys proteases of the herbivorous insect model Colorado potato beetle, *Leptinotarsa decemlineata*.^12^

Our goal in this study was to assess whether the non-functional elements of a cystatin, referred to collectively as the ‘scaffold’, would represent a relevant structural module for plant cystatin improvement. Cystatin scaffolds have been used to different ends in recent years, notably to develop accessory proteins useful in stabilizing and purifying recombinant proteins in heterologous protein expression systems,^13^ to design Affimer binding proteins for a variety of imaging, diagnostic and therapeutic purposes,^14,15^ or as an inspiration for *de novo* protein design involving customized protein folds.^16^ Here we used twelve cystatin scaffolds as supporting templates for the N-terminal trunk and inhibitory loops of alternative cystatins, to assess their usefulness in generating functional diversity among a finite collection of hybrid cystatin variants. The α-helix and inter-loop region of plant cystatins include positively selected amino acids presumably indicative of an impact on the protease inhibitory range or potency of the cystatin.^8,17^ The central fold of cystatins, although not directly involved in the inhibitory process, might represent a valuable target for plant cystatin engineering given its possible impact on the spatial orientation and stability of the protein’s functional elements.

## 2 RESULTS AND DISCUSSION

### 2.1 A generic scheme for cystatin scaffold substitution

A scaffold substitution strategy was designed (1) to assess whether grafting the N-terminal trunk and two inhibitory loops of a given cystatin on the scaffold of a second cystatin could be a viable option for the design of stable and active hybrid inhibitors, and (2) to determine whether natural cystatin scaffolds among the plant cystatin protein family could represent a pool of structural modules useful to generate cystatin variants with improved inhibitory potency against insect Cys proteases. A first set of hybrids were designed with the functional elements of tomato multicystatin domain SlCYS8, chosen based on the well described suitability of this inhibitor for protein engineering work^12,13,18,19^ and its moderate activity against arthropod Cys proteases, including *L. decemlineata* digestive Cys proteases, compared to other plant cystatins.^12^ A second set of hybrids were designed with the functional elements of *Physcomitrella patens* C-tailed cystatin domain PpCYS and potato multicystatin domain StCYS5, two cystatins highly active against herbivorous arthropod Cys proteases,^12^ to assess whether the inhibitory efficiency of naturally potent cystatins could be explained, at least in part, by their supporting scaffold. A third set of hybrids were designed with the N-terminal and inhibitory loops of cucumber cystatin CsCYS, a weak inhibitor of arthropod Cys proteases,^12^ to determine the relative influence of these functional elements and their supporting scaffold on the resulting potency of the cystatin.

Cystatin hybrids were designed *in silico* by ‘grafting’ the N-terminal trunk and two inhibitory loops of ‘donor’ cystatins onto the supporting scaffold of alternative, ‘recipient’ cystatins deemed representative of the plant cystatin family (**Table 1**). The three functional elements were defined based on their position relative to conserved amino acid motifs essential for protease inhibitory activity, in such a way as to include amino acids assumed to physically interact with amino acid residues in the active site or at the surface of the target enzyme (**Figure 1**).^9^ The scaffold corresponded to the polypeptide sequences remaining, including the α-helix and 1^st^ elbow between the N-terminal trunk and first inhibitory loop, the 2^nd^ elbow between the two inhibitory loops, and the amino acid string downstream of the second inhibitory loop at the C-terminal end. More specifically, the N-terminal trunks were defined based on the Gly–Gly (GG) motif conserved in the N-terminal trunk of plant cystatins, the first inhibitory loops based on the conserved pentapeptide motif Gln–X–Val–X–Gly (QxVxG) known to penetrate the active site of the target protease, and the second inhibitory loops based on the conserved Trp (W) residue also entering the active site cleft of the protease.^8^ Twenty hybrids were designed overall, with the N-terminal trunk and inhibitory loops of four donors (SlCYS8, PpCYS, StCYS5, CsCYS) and the supporting scaffolds of 12 recipients including the loop donors (**Table 2** and **Supplementary Table 1**). Synthetic DNA’s were generated for the 20 hybrids and used as templates for heterologous expression in *E. coli* and affinity purification using the GST gene fusion system.^20^ Eighteen hybrids, out of 20, were purified in a form suitable for protease inhibitory assays with papain and *L. decemlineata* Cys proteases (**Table 2** and **Figure 2**). The other two hybrids –with the functional elements of potato StCYS5– could not be properly expressed under our experimental conditions, likely due to unsuccessful folding and/or poor stability of the resulting protein products expressed in a foreign cellular environment.

**Table 1.**
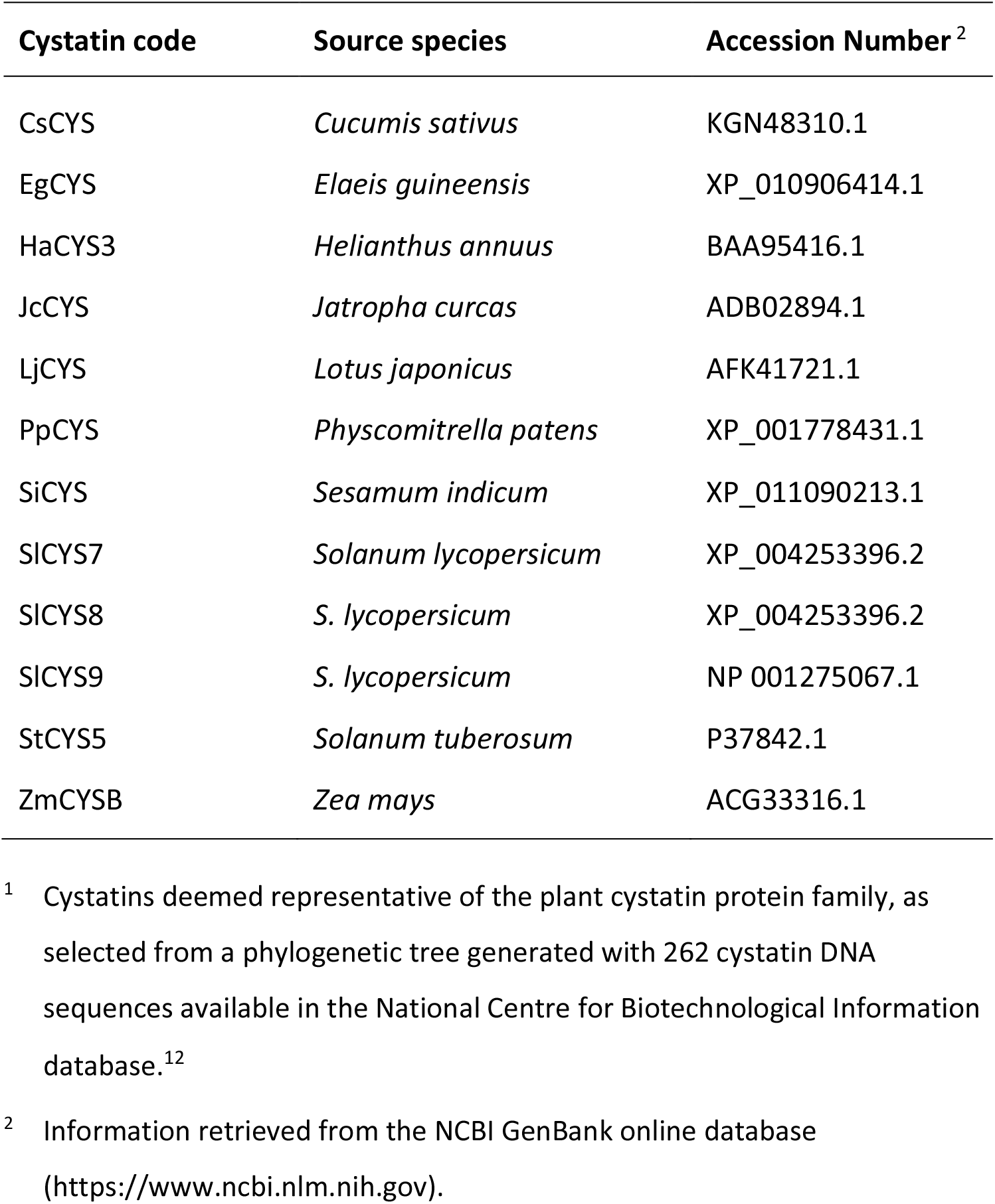
Plant cystatins selected for hybrid cystatin design ^1^

**FIGURE 1.**
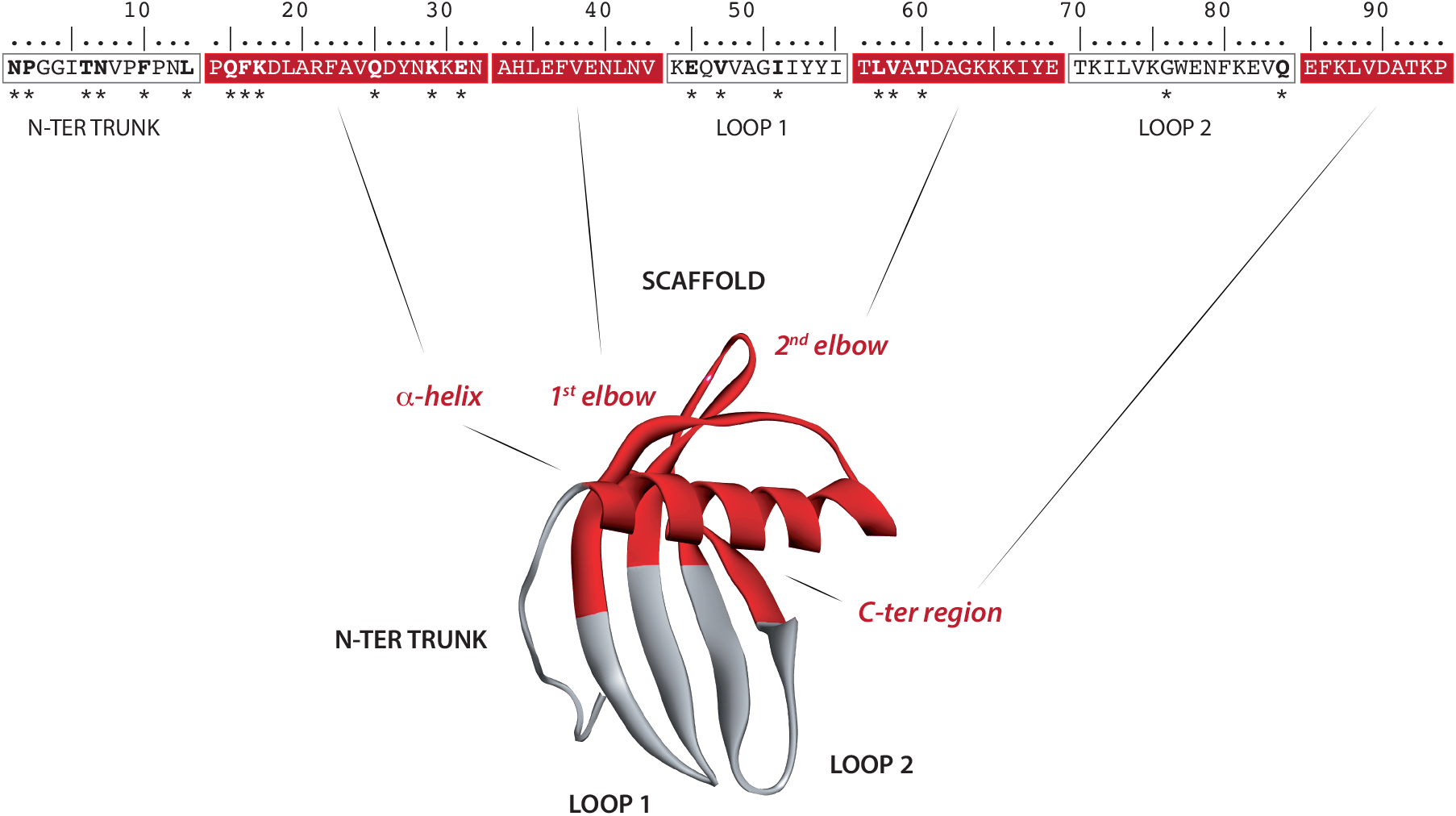
A generic scheme for plant cystatin scaffold substitution: tomato cystatin SlCYS8 as a working model. The cystatin hybrids were designed *in silico* by transferring the N-terminal trunk (N-ter trunk), first inhibitory loop (Loop 1) and second inhibitory loop (Loop 2) of a given cystatin on the supporting scaffold of an alternative cystatin. Genes synthesized for the resulting hybrids were then inserted in a modified pGEX-3X vector, downstream of a GST tag coding sequence for heterologous expression in *E. coli*. Polypeptide regions forming the scaffold of tomato SlCYS8 –α-helix, 1^st^ elbow, 2^nd^ elbow, C-terminal string– are shown in red. Asterisks (*) under the amino acid sequence highlight residues identified as being under positive selection in plant cystatins.^17^ The structural model for SlCYS8 was built using the Discovery Studio 2.5 protein modelling software (Accelrys, Inc.), based on the structural coordinates of oryzacystatin 1 (Protein Data Bank Accession 1EQK).

**Table 2.** Hybrid cystatins designed in this study, and their production rate following heterologous expression in *E. coli* and affinity purification using the GST gene fusion system

| Functional elements donor | Supporting scaffold (or recipient) <sup>1</sup> | Predicted mass (kDa) | Production rate <sup>2</sup> |
| --- | --- | --- | --- |
| SICYS8 | SICYS8 (ctrl) | 10.8 | Very high |
|  | CsCYS | 11.1 | Low |
|  | EgCYS | 10.9 | Very high |
|  | HaCYS3 | 10.3 | Very high |
|  | JcCYS | 11.5 | High |
|  | LjCYS | 10.9 | Very high |
|  | PpCYS | 10.9 | Very high |
|  | SiCYS | 10.2 | Very high |
|  | SICYS7 | 10.5 | Very high |
|  | SICYS9 | 10.7 | High |
|  | StCYS5 | 10.7 | High |
|  | ZmCYSB | 10.9 | Very high |
| CsCYS | CsCYS (ctrl) | 11.7 | Very high |
|  | PpCYS | 11.5 | Very high |
|  | SICYS8 | 11.3 | High |
|  | StCYS5 | 11.2 | Very high |
| PpCYS | PpCYS (ctrl) | 11.4 | Very high |
|  | CsCYS | 11.6 | High |
|  | SICYS8 | 11.3 | High |
|  | StCYS5 | 11.2 | Very high |
| StCYS5 | StCYS5 (ctrl) | 10.9 | Very high |
|  | CsCYS | 11.3 | None |
|  | PpCYS | 11.1 | Low |
|  | SICYS8 | 10.9 | None |

<sup>1</sup> See **Supplementary Table 1** for hybrids amino acid sequences.
<sup>2</sup> See **Figure 2** for selected examples resolved on gel.

**FIGURE 2.**
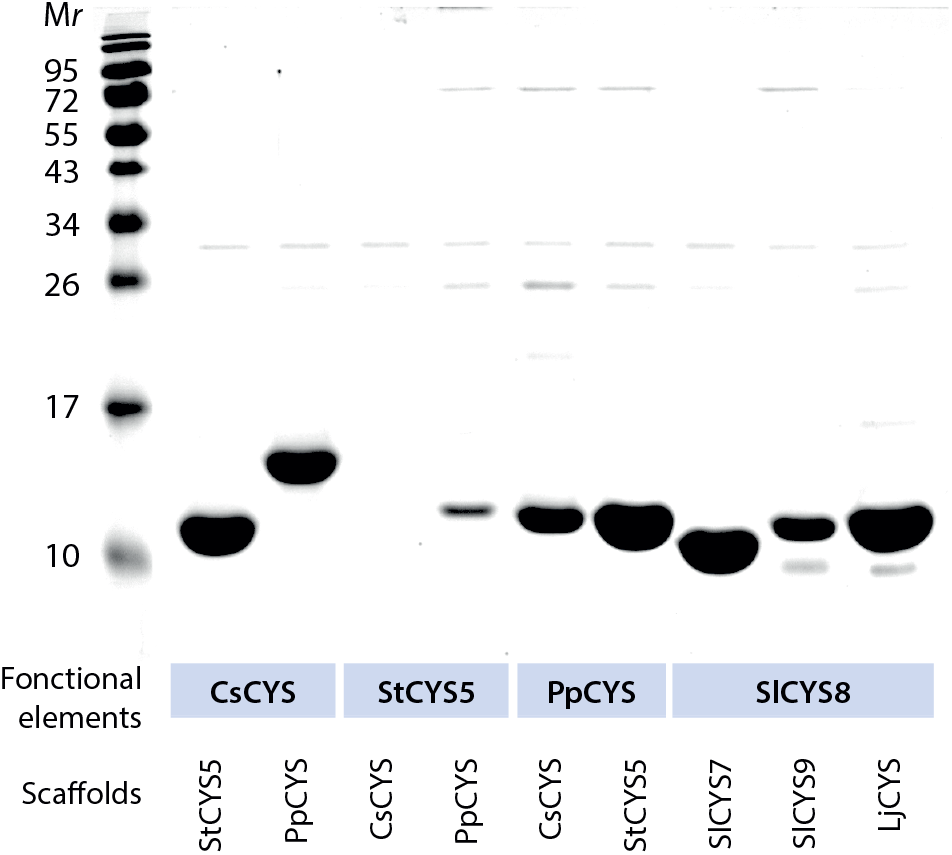
Selected subset of cystatin scaffold hybrids produced in this study, as visualized in Coomassie blue-stained polyacrylamide slab gels following 15% (w/v) SDS-PAGE in reducing conditions. Numbers on the left refer to the molecular mass of commercial protein markers. Details on the functional element donors and scaffold recipients are given in **Table 1, Table 2** and **Supplementary Table 1**.

### 2.2 The inhibitory potency of SlCYS8 functional elements is scaffold-dependent

Protease inhibitory assays were conducted to measure the impact of scaffold substitutions on the anti-papain activity of SlCYS8 N-terminal trunk and inhibitory loops (**Figure 3**). As recently observed with SlCYS8 hybrids bearing alternative inhibitory loops,^12^ *K*_i_ values for papain (*K*_i (papain)_ values) differed from one hybrid to another, positively or negatively depending on the scaffold. The most potent hybrids were those with the SlCYS8 functional elements grafted on the scaffolds of *Elaeis guineensis* cystatin EgCYS and potato StCYS5, which showed *K*_i (papain)_ values decreased by 2.1- and 1.7-fold, respectively, compared to wild-type SlCYS8. The less potent hybrids were those including the scaffolds of maize cystatin ZmCYSB and *Sesamum indicum* cystatin SiCYS, with *K*_i (papain)_ values increased by 2.8- and 1.7-fold, respectively, compared to SlCYS8. In several cases, the most potent scaffolds for this set of hybrids were from cystatins naturally exhibiting strong anti-papain activity. This was illustrated with EgCYS, StCYS5, tomato multicystatin domain SlCYS7 and *Jatropha curcas* cystatin JcCYS all exhibiting *K*_i (papain)_ values between two and six times lower than the corresponding value determined for wild-type SlCYS8.^12^ By contrast, no clear relation could be established between the negative impact of several other scaffolds on the activity of SlCYS8 functional elements and the anti-papain activity of the original cystatin recipients. For instance, the negative impacts observed above for the scaffolds of ZmCYSB and SiCYS could not be matched with *K*_i (papain)_ values for the wild-type versions of these two inhibitors, approximately three times lower than the *K*_i (papain)_ value determined for SlCYS8.^12^

**FIGURE 3.**
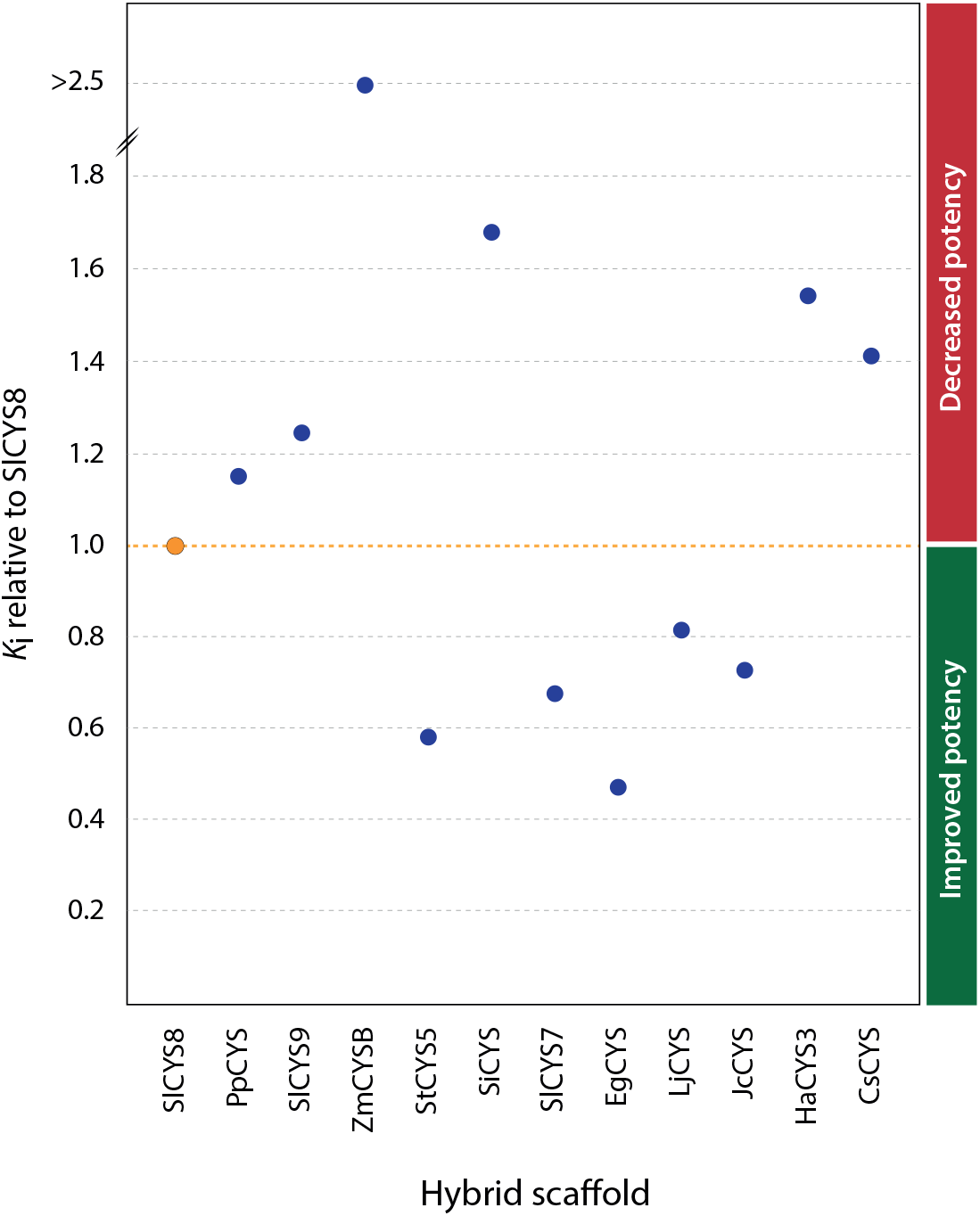
*K*_i (papain)_ values for the SlCYS8 scaffold hybrids, relative to the *K*_i (papain)_ value for wild-type SlCYS8 (ratio of 1.0). All hybrids designed in this study with the functional elements of SlCYS8 were produced in a stable form suitable for kinetic assays. A *K*_i (papain)_ ratio greater than 1.0 indicates a negative impact, and a *K*_i (papain)_ ratio lower than 1.0 a positive impact, of the recipient scaffold on papain inhibitory activity of the SlCYS8 functional elements.

Inhibitory assays were performed with the same set of hybrids to assess the potential of cystatin scaffold substitutions in generating hybrid inhibitors differentially active against digestive Cys cathepsins of *L. decemlineata* taken as an arthropod model (**Figure 4**). Several scaffold substitutions had an impact on the inhibitory potency of SlCYS8 functional elements against the insect cathepsin L-like enzymes (**Figure 4**, left panel). The most potent hybrids were those integrating the scaffolds of CsCYS, JcCYS and sunflower multicystatin domain HaCYS3, that showed relative inhibitory rates increased by >4 to 10-fold compared to wild-type SlCYS8 (*P* < 0.05; post-ANOVA Tukey’s test). The less potent hybrids were those with the scaffolds of PpCYS, SlCYS9, ZmCYSB and StCYS5, which had little effect on the inhibitory potency of SlCYS8 functional elements. Unlike for papain, no positive relation could be established between the basic efficiency of the original cystatins against *L. decemlineata* cathepsin L-like enzymes and the inhibitory potency of their resulting SlCYS8 hybrids against the same enzymes. For instance, the less potent hybrids included the scaffolds of cystatins, such as PpCYS, SlCYS9 or StCYS5, exhibiting strong inhibitory activity against the insect proteases.^12^ In sharp contrast, the most potent inhibitors, with the scaffolds of CsCYS, HaCYS3, JcCYS or lotus cystatin LjCYSB, were derived from cystatins exhibiting low to very weak activity against these enzymes.^12^

**FIGURE 4.**
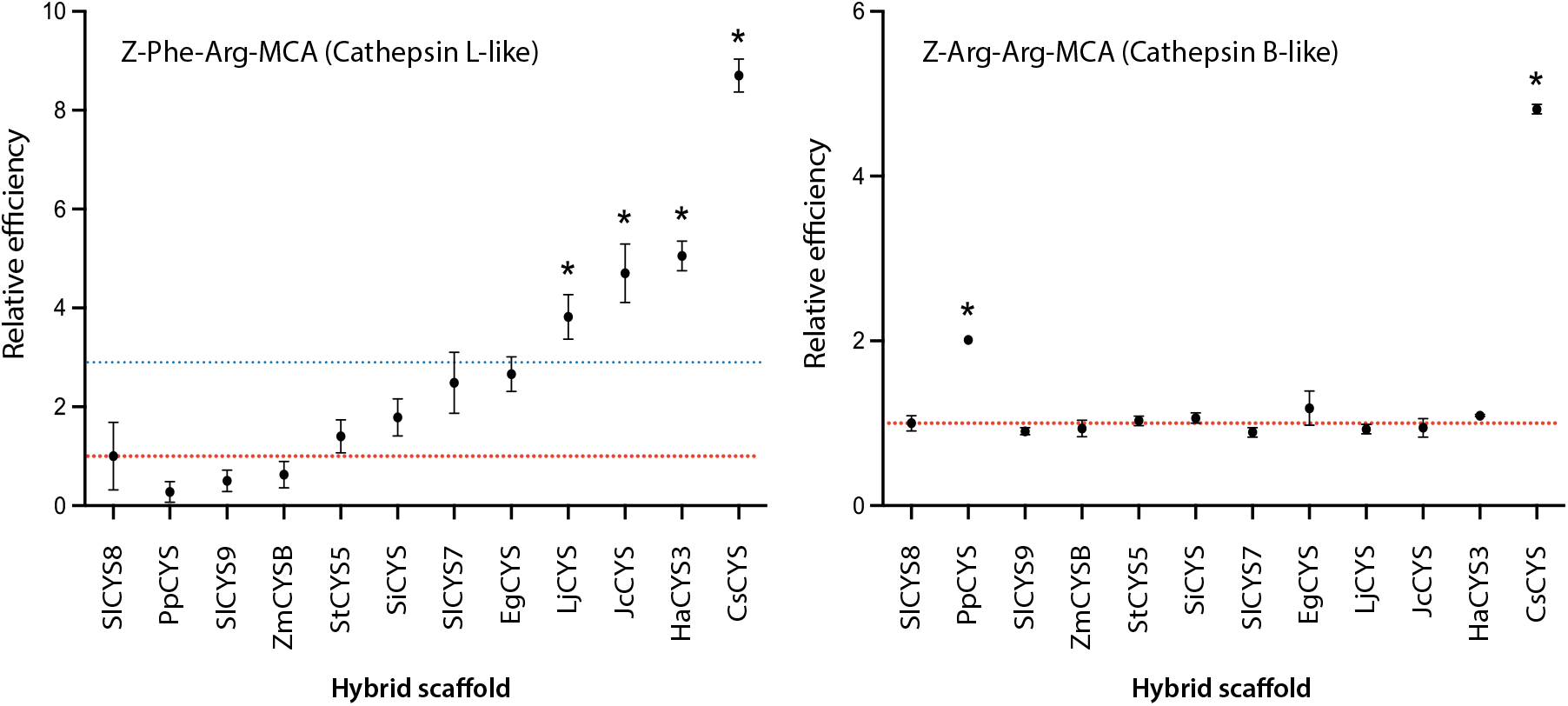
Inhibition of *L. decemlineata Z*-Phe–Arg-MCA-hydrolyzing (cathepsin L-like) and *Z*-Arg–Arg-MCA-hydrolyzing (cathepsin B-like) digestive proteases by the SlCYS8 scaffold hybrids. Data are expressed as relative inhibitory efficiency rates relative to the efficiency rate of wild-type SlCYS8 (ratio of 1; red dotted line). An inhibitory rate greater than 1.0 indicates a positive impact, and an inhibitory rate lower than 1.0 a negative impact, of the scaffold on inhibitory activity of the SlCYS8 functional elements. The inhibitory assays were conducted with cystatin concentrations of 250 nM [cathepsin L activity] or 1 µM [cathepsin B activity] in the reaction mixture. Each datapoint is the mean of three independent replicates ± SE. Asterisks indicate significantly different values compared to SlCYS8 (post-ANOVA Dunnett’s test, with an α value threshold of 5%). The blue line on left panel highlights average inhibitory efficiency of the 11 scaffold hybrids against cathepsin L-like enzymes, 2.9-fold the efficiency of wild-type SlCYS8.

Complementary assays were performed to assess the inhibitory potency of SlCYS8 hybrids against *L. decemlineata* cathepsin B-like enzymes (**Figure 4**, right panel). In accordance with previous studies reporting the poor susceptibility of these enzymes to plant cystatin inhibition,^10,20^ most scaffold substitutions had no effect on the effectiveness of SlCYS8 functional elements to inhibit their activity. Unexpectedly, the scaffolds of PpCYS, a strong inhibitor of herbivorous arthropod Cys proteases,^12^ and CsCYS, a weak inhibitor of these enzymes, had a positive impact on the SlCYS8 elements against these ‘naturally insensitive’ proteases, as inferred from inhibitory rates increased by 2- to 5-fold compared to wild-type SlCYS8. These observations, along with those above about cathepsin L-like enzymes, suggested a role for the supporting scaffold on the inhibitory potency of plant cystatin functional elements against Cys proteases. In practice, they suggested the potential of plant cystatin scaffolds as structural modules to generate functional diversity among small collections of recombinant cystatin variants, in complement to current protein engineering approaches relying on primary sequence modifications in the N-terminal trunk and two inhibitory loops.

### 2.3 The scaffold is a strong determinant of plant cystatins inhibitory potency

Cystatin hybrids designed with the N-terminal trunk and inhibitory loops of PpCYS, StCYS5 and CsCYS were used to assess the relative contributions of their scaffold to the inhibition of *L. decemlineata* proteases (**Figure 5**). In brief, cystatin hybrids with the functional elements of PpCYS or StCYS5 grafted onto alternative scaffolds showed decreased potency against the insect cathepsin L-like enzymes compared to their wild-type counterpart (**Figure 5**, upper panel). As expected, given the poor efficiency of CsCYS functional elements against *L. decemlineata* Cys proteases,^12^ CsCYS hybrids showed very low inhibitory activity against the same enzymes. Similar trends were observed for the insect cathepsin B-like enzymes (**Figure 5**, lower panel), with wild-type PpCYS and StCYS5 showing stronger activity than their resulting hybrids on alternative scaffolds, and CsCYS hybrids showing barely detectable activity against these enzymes as also observed with wild-type CsCYS. Considering the negligible impact of scaffold substitutions on the inhibitory potency of CsCYS bearing weakly active functional elements, the significant loss of inhibitory efficiency for the functional elements of PpCYS and StCYS5 grafted onto an alternative scaffold, and the conclusions drawn above with the SlCYS8 hybrids, our observations indicated overall an important impact of both the function-related structural elements and supporting scaffold on the inhibitory potency of plant cystatins.

**FIGURE 5.**
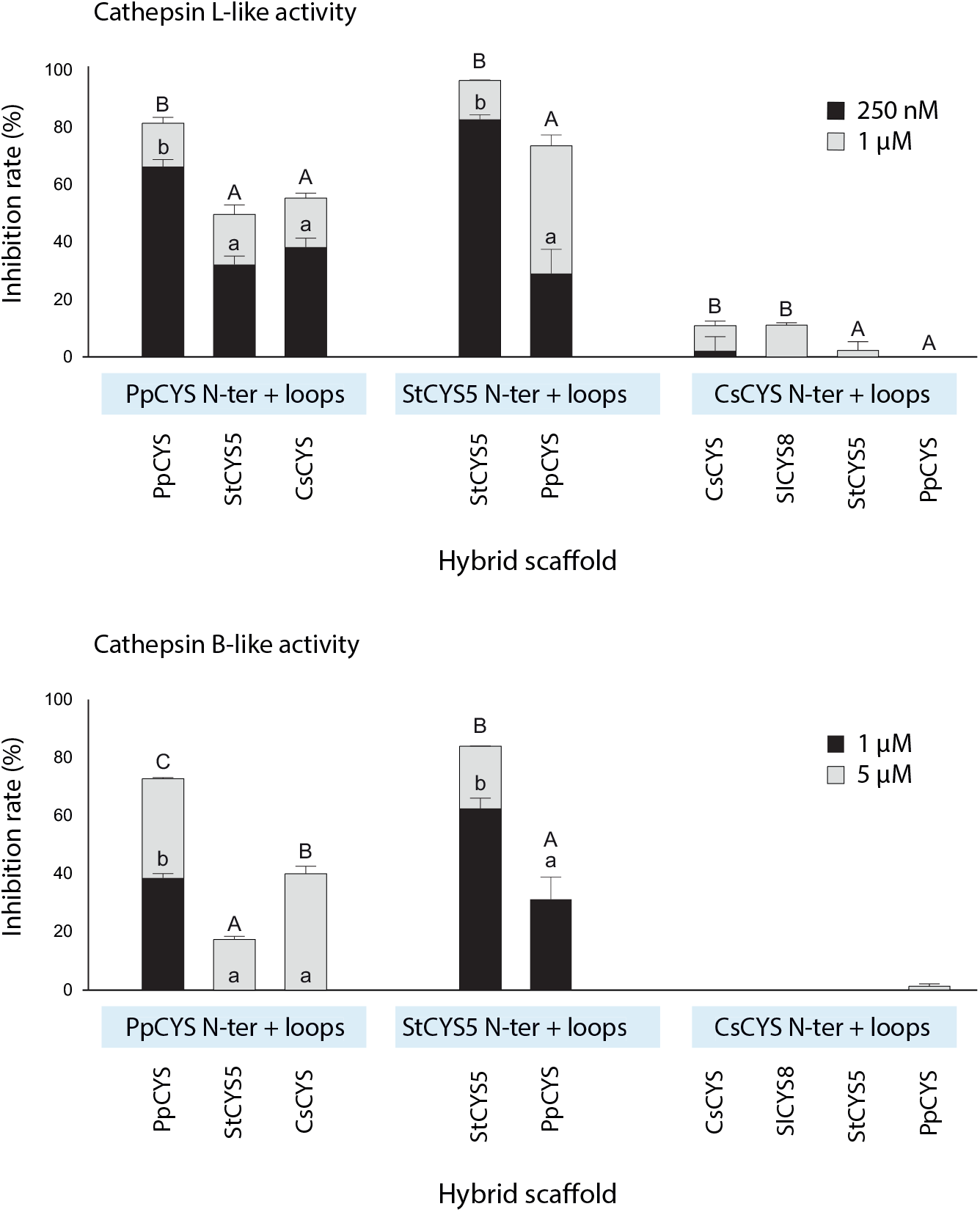
Inhibition of *L. decemlineata Z*-Phe–Arg-MCA-hydrolyzing (cathepsin L) and *Z*-Arg–Arg-MCA-hydrolyzing (cathepsin B) enzymes by cystatin hybrids with the functional elements of PpCYS, StCYS5 or CsCYS grafted onto alternative scaffolds. Data are expressed as relative inhibitory rates compared to the inhibitory rate measured with cysteine protease diagnostic inhibitor E-64 (arbitrary value of 100%). Inhibitory assays were conducted with limiting (250 nM [cathepsin L] or 1 µM [cathepsin B]) or excess (1 µM [cathepsin L] or 5 µM [cathepsin B]) concentrations of cystatin. Each bar is the mean of three independent replicates ± SD. For each set of functional elements, different letters (lower case for limiting concentrations, capital letters for excess concentrations) indicate significantly different inhibitory rates among the cystatin hybrids (post-ANOVA Tukey’s tests, with an α value threshold of 5%).

Distribution maps were inferred from the *L. decemlineata* cathepsin L inhibition data to compare the potential of our new scaffold substitution strategy in generating functional variability with our recently described strategy based on the substitution of N-terminal trunks or inhibitory loops (**Figure 6**). An overall coefficient of variation (CV) of 62% was calculated for inhibitory data generated with the scaffold hybrids, similar to a CV value of 61% calculated for the cathepsin L inhibition data recently produced with SlCYS8 structural element (SE) hybrids.^12^ Much interestingly, the average cathepsin L inhibitory rate for the SlCYS8 scaffold hybrids was almost three times the inhibitory rate measured for wild-type SlCYS8 (*see* **Figure 4**), similar to the improved inhibitory rates observed for the best variants of a previously described collection of SlCYS8 single mutants bearing alternative residues at positively selected sites.^20^ Together, these observations confirmed the potential of our new scaffold substitution approach to implement functional diversity among a small set of recombinant cystatin variants considered for plant protection, in complement to the usual mutational approaches involving structural changes in the functional elements.

**FIGURE 6.**
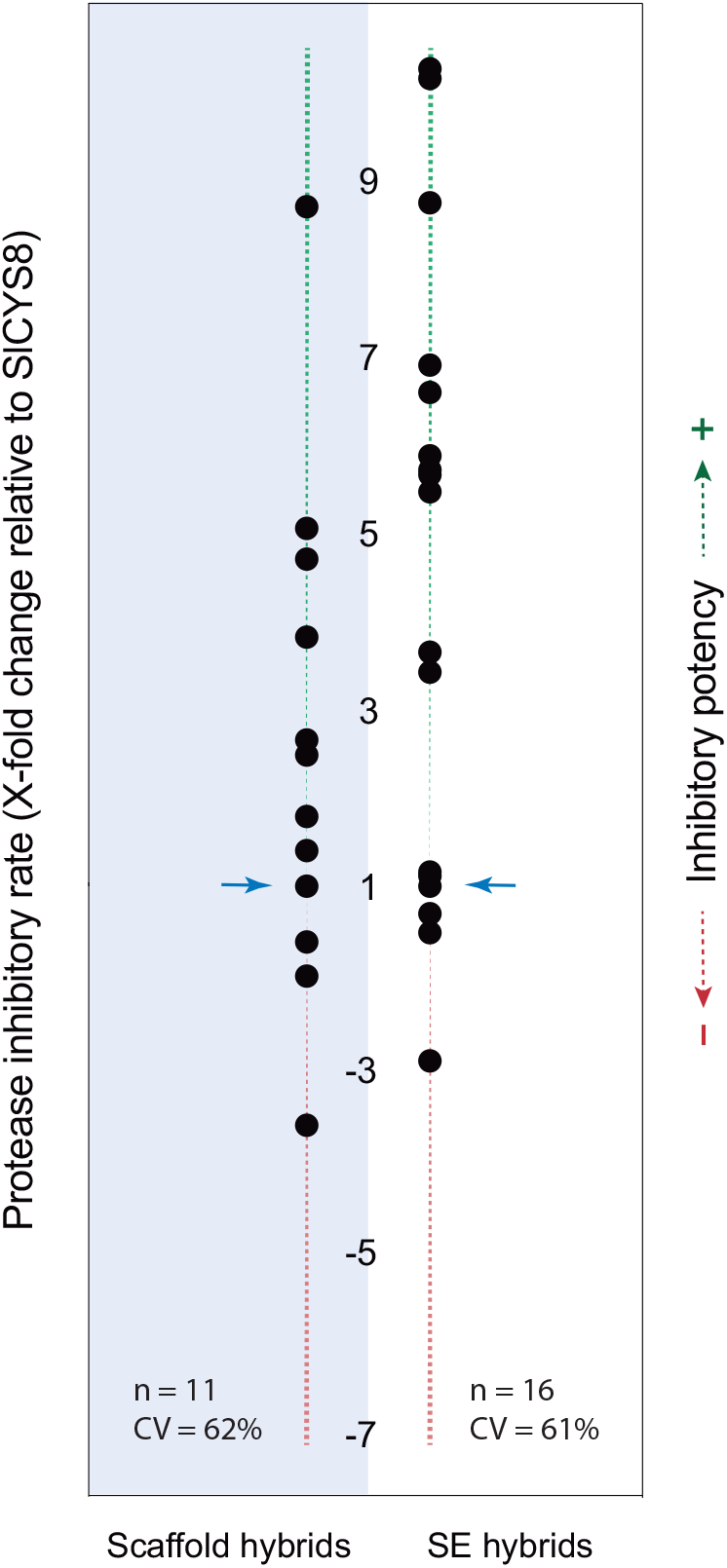
Functional variation among finite collections of SlCYS8 hybrids bearing an alternative scaffold [Scaffold hybrids] (**Figure 4**, this study) and SlCYS8 hybrids bearing the function-related structural elements (SE’s) of an alternative cystatin [SE hybrids] (Figure 6 of ref. 12). Data are expressed as *L. decemlineata* cathepsin L inhibitory rates for the SlCYS8 hybrids at limiting concentration, relative to the inhibitory rate determined for wild-type SlCYS8. Blue arrows point to wild-type SlCSY8, with a relative inhibitory rate of 1. n, number of hybrids considered; CV, coefficients of variation, as determined with the absolute values of X-fold change ratios.

## 3 CONCLUSION

Protein engineering approaches have been proposed to improve the inhibitory properties of plant cystatins against herbivorous arthropod digestive Cys proteases, typically involving rationally inferred or randomly generated mutations at functionally relevant amino acid sites of the protein.^2,11^ We recently described a novel approach for cystatin engineering, where the N-terminal trunk and/or inhibitory loops of a given cystatin are changed for the corresponding element(s) of an alternative cystatin.^12,21^ Here, we assessed whether the scaffold region supporting these functional elements could also represent a valuable structural module for cystatin function improvement. Hybrid cystatins were designed *in silico*, consisting of the N-terminal trunk and two inhibitory loops of a given donor cystatin grafted onto the scaffold of an alternative recipient cystatin, and then tested for their inhibitory potency against papain and digestive Cys cathepsins of *L. decemlineata* used as a plant pest model. The scaffold region of plant cystatins includes hypervariable, positively selected amino acid sites thought to influence the inhibitory activity of the protein against Cys proteases.^17^ Supporting this, we showed previously single SlCYS8 mutants bearing alternative amino acids at position 31 in the α-helix, in place of the original glutamate residue, to exhibit improved activity against several Cys proteases including *L. decemlineata* cathepsin L-like enzymes.^20^ Accordingly, our data in this study confirmed a significant impact of the supporting scaffold on the inhibitory activity of SlCYS8 and other plant cystatins. In practice, they suggested the potential of plant cystatin scaffolds as useful structural modules to design recombinant cystatins with improved activity against the digestive Cys proteases of herbivorous arthropods.

Studies will be welcome in coming years to compare the plant protective effects of our most potent SlCYS8 scaffold hybrids with the protective effects of recently described SlCYS8 SE hybrids^12^ or those of ^P2V^SlCYS8, a single variant of SlCYS8 reported to compromise *L. decemlineata* larval growth.^5^ Studies will also be welcome to assess pleiotropic effects of the scaffold hybrids *in planta*, considering the possible off-target effects of recombinant cystatins in transgenic plant lines developed for pest control.^22^ Agronomically valuable phenotypes have been described in plants modified to express recombinant cystatins,^22-27^ but negative effects impacting Cys protease-dependent cellular processes or compromising plantlet regeneration following leaf cell genetic transformation cannot be ruled out at this stage.^2,28^

## 4 MATERIALS AND METHODS

### 4.1 Recombinant cystatins

All cystatins were produced in *E. coli*, strain BL21 as described previously,^20^ using the GST gene fusion system for heterologous expression and affinity purification (GE Healthcare). DNA templates for the cystatins were synthesized as g-blocks (Integrated DNA Technologies) including GoldenGate BsaI cloning sites on both sides of the cystatin coding region. DNA coding sequences for the original cystatins corresponded to those reported in GenBank (as listed in **Table 1**). DNA sequences for the scaffold hybrids were designed as described on **Figure 1**, with the functional elements of donor cystatins SlCYS8, StCYS5, PpCYS or CsCYS grafted onto the cystatin scaffold recipients as detailed in **Supplementary Table 1**. The genes were inserted in a modified version of the pGEX-3X expression vector (GE Healthcare) using the Golden Gate DNA shuffling method of Engler et al.,^29^ downstream of a ‘GST–factor Xa cleavage site’ coding sequence.^30^ The GST tag was removed by cleavage with bovine factor Xa, according to the manufacturer’s specifications (Millipore Sigma). The purified cystatins were quantified by densitometric analysis of Coomassie blue-stained polyacrylamide slab gels following 15% (w/v) SDS-PAGE, using three technical replicates and bovine serum albumin (Sigma-Aldrich) as a protein standard.

### 4.2 Test proteases

Papain (E.C.3.4.22.2, from papaya latex) was from Millipore Sigma. Insect proteases were extracted from the midgut of *L. decemlineata* fourth instars collected on potato plants, as described previously.^20^ Papain and soluble proteins in the arthropod crude extracts were assayed according to Bradford,^31^ with bovine serum albumin as a protein standard.

### 4.3 *K*_i (papain)_ value determinations

*K*_i (papain)_ values for the cystatin hybrids were determined by the monitoring of substrate hydrolysis progress curves based on the linear equation of Henderon.^32,33^ Papain activity was monitored in 50 mM Tris-HCl, pH 7.0, using the synthetic peptide substrate *Z*-Phe–Arg-methylcoumarin (MCA) (Sigma-Aldrich). Hydrolysis was allowed to proceed at 25°C in reducing conditions (10 mM L-cysteine) with the substrate in large excess, after adding (or not) recombinant cystatins diluted in a minimal volume of reaction buffer. Papain activity was monitored using a Synergy H1 fluorimeter (BioTek), under an excitation wavelength of 360 nm and an emission wavelength of 450 nm. *K*_i_ values were calculated with experimentally determined *K*_i(app)_ and *K*_m_ values, using the following equation: *K*_i_ = *K*_i(app)_ / (1 + [S] / *K*_m_). A *K*_m_ value of 93.6 µM was determined for papain under our assay conditions.

### 4.4 Arthropod protease assays

Cys cathepsin activities in *L. decemlineata* protein extracts were assayed in 0.1 M NaH_2_PO_4_/0.1 M Na_2_HPO_4_ phosphate buffer, pH 6.5, using the synthetic peptide substrates *Z*-Phe–Arg-MCA for cathepsin L-like and *Z*-Arg–Arg-MCA for cathepsin B-like activities.^12^ Hydrolysis was allowed to proceed in reducing conditions (10 mM L-cysteine) for 10 min at 25°C, with ∼5 ng of arthropod protein per µl in the reaction mixture and the peptide substrate in large excess. Cystatins diluted in a minimal volume of reaction buffer were added to the reaction mixture for the inhibitory assays. Proteolytic activity was monitored using a Synergy H1 fluorimeter (BioTek), under excitation and emission wavelengths of 360 nm and 450 nm, respectively.

## AUTHOR CONTRIBUTIONS

JT, CG and DM conceived the protein design approach and experimental strategy. JT and MCG performed the experiments and analyzed the data. JT, MCG and DM wrote the manuscript. All authors read and approved the manuscript for publication.

## SUPPORTING INFORMATION

Additional supporting information may be found online in the Supporting Information section at the end of this article.

## ACKNOWLEDGMENTS

This work was supported by a Discovery grant from the Natural Science and Engineering Research Council of Canada (RGPIN-2015-06022).

## Abbreviations

CsCYS: *Cucumis sativa* (cucumber) cystatin
CV: coefficient of variation
GST: glutathione *S*-transferase
MCA: methylcoumarin
SlCYS8: eighth inhibitory domain of tomato multicystatin
StCYS5: fifth inhibitory domain of potato multicystatin
PpCYS: *Physcomitrella patens* cystatin

