## Supplementary Table 1 for "Hidden potential of the supporting scaffold as a structural module for plant cystatin design"

Centre de recherche et d'innovation sur les végétaux, Université Laval

SUPPORTING INFORMATION

**Supplementary Table 1** Amino acid sequences of tomato SlCYS8, potato StCYS5, *Physcomitrella patens* PpCYS and cucumber CsCYS scaffold hybrids designed in this study.

**Supplementary Table 1** Amino acid sequences of the tomato SlCYS8, potato StCYS5, *Physcomitrella patens* PpCYS and cucumber CsCYS scaffold hybrids designed in this study. Polypeptide segments of the supporting scaffolds are shown in *italics*.

| Loops | Scaffold | N-terminal trunk | α-helix + 1st elbow | Loop 1 | 2nd elbow | Loop 2 | C-terminal string |
|---|---|---|---|---|---|---|---|
| SlCYS8 | SlCYS8 (ctrl) | 1–13 NPGGITNVPFPNL | *14–43 PQFKDLARFAVQDYNKKENAHLEFVENLNV* | 44–55 KEQVVAGIIYYI | *56–69 TLVATDAGKKKIYE* | 70–84 TKILVKGWENFKEVQ | *85–95 EFKLVGDATKP* |
| | CsCYS | 1–13 NPGGITNVPFPNL | *14–43 SRMKELAEWAVAEHNKKAGTHLMFIGILTC* | 44–55 KEQVVAGIIYYI | *56–72 TLTAKDEKDNCEIESYM* | 73–87 TKILVKGWENFKEVQ | *88–97 YFQKLLLAEQ* |
| | EgCYS | 1–13 NPGGITNVPFPNL | *14–43 VEIEELARFAVEEHNKKANTLLEFGRVVKV* | 44–55 KEQVVAGIIYYI | *56–69 TIEASDGGKKKLYE* | 70–84 TKILVKGWENFKEVQ | *85–97 EFIPLGDCSSENE* |
| | HaCYS3 | 1–13 NPGGITNVPFPNL | *14–43 LVIDDLARFAVDEYSKKQNTLLEFERVLDA* | 44–55 KEQVVAGIIYYI | *56–69 ILEATVGGVKNTYV* | 70–84 TKILVKGWENFKEVQ | *85–90 EFKHLY* |
| | JcCYS | 1–13 NPGGITNVPFPNL | *14–43 VEIDGLARFAVDDYNKKQNALLEYKRVVNA* | 44–55 KEQVVAGIIYYI | *56–69 TLEVEDGGQKKVYE* | 70–84 TKILVKGWENFKEVQ | *85–103 EFKLIGDAPTQSTPTESTA* |
| | LjCYSB | 1–13 NPGGITNVPFPNL | *14–43 LEIDALARFAVDEHNNKQNALLEFGRVVKA* | 44–55 KEQVVAGIIYYI | *56–69 TLEAIDGGEKKIYE* | 70–84 TKILVKGWENFKEVQ | *85–98 EFKEVAGDAPLFTT* |
| | PpCYS | 1–13 NPGGITNVPFPNL | *14–45 LEIDEAAKFAVAEHNDRENSLEKLTFSKVVSC* | 46–57 KEQVVAGIIYYI | *58–71 VIEVEEGSSIKLYE* | 72–86 TKILVKGWENFKEVQ | *87–97 EFKLKDAGVTSA* |
| | SiCYS | 1–13 NPGGITNVPFPNL | *14–43 TEINDLARFAVNEQQKKENTLLEFKKVLSA* | 44–55 EQVVAGIIYYI | *56–69 TLEAVEGGKSKVYE* | 70–84 TKILVKGWENFKEVQ | *85–92 EFKLIGDA* |
| | SlCYS7 | 1–13 NPGGITNVPFPNL | *14–43 PPLQDLARFAVQDYNKAQNAHLEYVENLNV* | 44–55 KEQVVAGIIYYI | *56–69 TLVATDAGKKKIYE* | 70–84 TKILVKGWENFKEVQ | *85–93 EFKLVGDDS* |
| | SlCYS9 | 1–13 NPGGITNVPFPNL | *14–43 DEIHSLAKFAVDEHNKKENAMIELARVVKA* | 44–55 KEQVVAGIIYYI | *56–69 TLEVMDAGKKKLYE* | 70–84 TKILVKGWENFKEVQ | *85–94 EFKHVEDVPT* |

**Supplementary Table 1** Cont'd

| Loops | Scaffold | N-terminal trunk | α-helix + 1st elbow | Loop 1 | 2nd elbow | Loop 2 | C-terminal string |
|---|---|---|---|---|---|---|---|
| SlCYS8 | StCYS5 | 1–13<br>NPGGITNVPFPNL | *14–43*<br>*PEFQDLTRFAVHQYNKDQNAHLEFVENLNV* | 44–55<br>KEQVVAGIIYYI | *56–69*<br>*TFAATDGGKKKIYE* | 70–84<br>TKILVKGWENFKEVQ | *85–94*<br>*EFKLVGDDSA* |
| | ZmCYSB | 1–13<br>NPGGITNVPFPNL | *14–43*<br>*AESDGLGRFAVDEHNRRENALLEFVRVVEA* | 44–55<br>KEQVVAGIIYYI | *56–69*<br>*TLEAVEAGRKKLYE* | 70–84<br>TKILVKGWENFKEVQ | *85–97*<br>*EFSHKGDATAFTN* |
| CsCYS | CsCYS<br>(ctrl) | 1–18<br>MASDLVPGGYTPVENPQS | *19–48*<br>*SRMKELAEWAVAEHNKKAGTHLMFIGILTC* | 49–60<br>ESQIVDGVNYRF | *61–77*<br>*TLTAKDEKDNCEIESYM* | 78–92<br>AVVFEQPWEHIKELV | *93–102*<br>*YFQKLLLAEQ* |
| | PpCYS | 1–18<br>MASDLVPGGYTPVENPQS | *19–50*<br>*LEIDEAAKFAVAEHNDRENSLEKLTFSKVVSC* | 51–62<br>ESQIVDGVNYRF | *63–76*<br>*VIEVEEGSSIKLYE* | 77–92<br>AVVFEQPWEHIKELV | *93–104*<br>*EFKLKDAGVTSA* |
| | SlCYS8 | 1–18<br>MASDLVPGGYTPVENPQS | *19–48*<br>*PQFKDLARFAVQDYNKKENAHLEFVENLNV* | 49–60<br>ESQIVDGVNYRF | *61–74*<br>*TLVATDAGKKKIYE* | 75–89<br>AVVFEQPWEHIKELV | *90–100*<br>*EFKLVGDATKP* |
| | StCYS5 | 1–18<br>MASDLVPGGYTPVENPQS | *19 –48*<br>*PEFQDLTRFAVHQYNKDQNAHLEFVENLNV* | 49 –60<br>ESQIVDGVNYRF | 61 –74<br>*TFAATDGGKKKIYE* | 75 –89<br>AVVFEQPWEHIKELV | *90 –99*<br>*EFKLVGDDSA* |
| PpCYS | PpCYS<br>(ctrl) | 1–16<br>MLSGGKQEVDLQNSNN | *17–48*<br>*LEIDEAAKFAVAEHNDRENSLEKLTFSKVVSC* | 49–60<br>HMQVVAGSMYYL | *61–74*<br>*VIEVEEGSSIKLYE* | 75–89<br>AKVWVKPWQNFKKLE | *90–101*<br>*EFKLKDAGVTSA* |
| | CsCYS | 1–16<br>MLSGGKQEVDLQNSNN | *17–46*<br>*SRMKELAEWAVAEHNKKAGTHLMFIGILTC* | 47–58<br>HMQVVAGSMYYL | *59–74*<br>*TLTAKDEKDNCEIESYM* | 75–89<br>AKVWVKPWQNFKKLE | *90–99*<br>*YFQKLLLAEQ* |
| | SlCYS8 | 1–16<br>MLSGGKQEVDLQNSNN | *17–46*<br>*PQFKDLARFAVQDYNKKENAHLEFVENLNV* | 47–58<br>HMQVVAGSMYYL | *59–72*<br>*TLVATDAGKKKIYE* | 73–87<br>AKVWVKPWQNFKKLE | *88–98*<br>*EFKLVGDATKP* |
| | StCYS5 | 1–16<br>MLSGGKQEVDLQNSNN | *17–46*<br>*PEFQDLTRFAVHQYNKDQNAHLEFVENLNV* | 47–58<br>HMQVVAGSMYYL | *59–72*<br>*TFAATDGGKKKIYE* | 73–87<br>AKVWVKPWQNFKKLE | *88–97*<br>*EFKLVGDDSA* |

**Supplementary Table 1** Cont'd

| Loops | Scaffold | N-terminal trunk | α-helix + 1st elbow | Loop 1 | 2nd elbow | Loop 2 | C-terminal string |
|---|---|---|---|---|---|---|---|
| StCYS5 | StCYS5 (ctrl) | 1–13<br>KLGGFTEVPFPNS | *14–43*<br>*PEFQDLTRFAVHQYNKDQNAHLEFVENLNV* | 44–55<br>KKQVVAGMLYYI | *56–69*<br>*TFAATDGGKKKIYE* | 70–84<br>TKIWVKVWENFKKVV | *85–94*<br>*EFKLVGDDSA* |
| | CsCYS | 1–13<br>KLGGFTEVPFPNS | *14–43*<br>*SRMKELAEWAVAEHNKKAGTHLMFIGILTC* | 44–55<br>KKQVVAGMLYYI | *56–72*<br>*TLTAKDEKDNCEIESYM* | 73–87<br>TKIWVKVWENFKKVV | *88–97*<br>*YFQKLLLAEQ* |
| | PpCYS | 1–13<br>KLGGFTEVPFPNS | *14–45*<br>*LEIDEAAKFAVAEHNDRENSLEKLTFSKVVSC* | 46–57<br>KKQVVAGMLYYI | *58–71*<br>*VIEVEEGSSIKLYE* | 72–86<br>TKIWVKVWENFKKVV | *87–98*<br>*EFKLKDAGVTSA* |
| | SlCYS8 | 1–13<br>KLGGFTEVPFPNS | *14–43*<br>*PQFKDLARFAVQDYNKKENAHLEFVENLNV* | 44–55<br>KKQVVAGMLYYI | *56–69*<br>*TLVATDAGKKKIYE* | 70–84<br>TKIWVKVWENFKKVV | *85–95*<br>*EFKLVGDATKP* |
